# Macrophages allocate independent of apoptosis and produce reactive oxygen species during interdigital phagocytosis

**DOI:** 10.1101/2023.05.04.539494

**Authors:** David Hernández-García, Celina García-Meléndrez, Rocío Hernández-Martínez, Omar Collazo-Navarrete, Luis Covarrubias

## Abstract

During programmed cell death (PCD), it is commonly accepted that macrophages are recruited by apoptotic cells to complete cell degradation. Interdigital cell death, a classical model of PCD, contributes to digit individualization in limbs of mammals and other vertebrates. Here we show that macrophages are present in interdigits before significant cell death occurs and remain after apoptosis inhibition. The typical interdigital phagocytic activity was not observed after a partial depletion of macrophages and was markedly reduced by engulfment/phagosome maturation inhibition, as detected by its association with high lysosomal activity. β-galactosidase activity in this region was also coupled with phagocytosis, against its relationship with cellular senescence. Interdigital phagocytosis correlated with high levels of reactive oxygen species (ROS), common in embryo regions carrying abundant cell death, suggesting that macrophages are the major source of ROS. ROS generation was dependent on NADPH oxidases and blood vessel integrity, but not directly associated with lysosomal activity. Therefore, macrophages prepattern regions where abundant cell death is going to occur, and high lysosomal activity and the generation of ROS by an oxidative burst-like phenomenon are activities of phagocytosis.

**Summary statement:** Recruitment of macrophages to the interdigital regions is not linked to apoptosis initiation and they phagocytize by a mechanism involving high lysosomal activity and an oxidative burst-like phenomenon.

## INTRODUCTION

Programmed cell death (PCD) plays a fundamental role in embryo development and regulates tissue homeostasis (Fuchs and Steller, 2011). Although different types of PCD exist (Bedoui et al., 2020), apoptosis is the best characterized form of PCD occurring in animal development (Conradt, 2009). The regulation of apoptosis is complex but once initiated cells display stereotyped morphological and molecular changes. Changes in cell morphology include plasma membrane blebbing, cell shrinkage, chromatin condensation and nuclear fragmentation, whereas from the molecular point of view, apoptosis is executed by caspases which cut a variety of substrates causing a rapid and ordered cell degradation (Green and Llambi, 2015; Nicholson, 1999; Voss and Strasser, 2020). Generally, the efficient clearance of apoptotic bodies occurs within phagocytes either professional (macrophages and dendritic cells) or non-professional (epithelial and fibroblast cells) (Medina and Ravichandran, 2016).

Phagocytosis of apoptotic cells is fundamental for a successful apoptosis. Clearance of apoptotic cells by this process is assumed to prevent the release of intracellular components that could potentially lead to inflammation and damage to the surrounding tissue. Phagocytic cells, mainly macrophages in mammals, can be recruited by apoptotic cells through the release of chemotactic molecules (“find-me” signals) (Lemke, 2019; Medina and Ravichandran, 2016), though none has been identified in association with developmental cell death. Cells undergoing apoptosis expose signals to the external face of plasma membrane (“eat-me” signals; e.g., phosphatidylserine– PS) that are recognized by phagocytes for cell engulfment (Lemke, 2019; Nagata, 2018). Most of the apoptotic bodies *in vivo* are found within macrophages suggesting the promptness and high efficiency of the engulfment process (Nagata, 2018; Rotello et al., 1994; Wood et al., 2000).

The above description establishes macrophage recruitment and the subsequent engulfment as processes occurring late in the course of apoptosis (Poon et al., 2010). In vivo, this phenomenon has been observed during cochlear development, in which case macrophages are recruited to the regressing greater epithelial ridge after apoptosis initiation (Borse et al., 2021). However, macrophages appear to be required for the activation of cell death during hyaloid vessel remodeling in the developing eye (Lobov et al., 2005) and for the natural death of Purkinje and motor neurons (Marin-Teva et al., 2004; Sedel et al., 2004), scenarios in which macrophages should be recruited before apoptosis initiation. Although the allocation of the so called resident macrophages has been studied (Perdiguero and Geissmann, 2016; Stremmel et al., 2018) and macrophages in regions undergoing abundant cell death have been found (Henson and Hume, 2006; Wood and Martin, 2017), the specific mechanisms involved in macrophage recruitment during development are largely unknown.

Interdigital cell death (ICD) contributes to digit individualization by controlling growth and sculpting the limb tissue (Hernández-Martínez and Covarrubias, 2011; Lorda-Diez et al., 2015a). Most of the cell death in this region occurs by caspase-dependent apoptosis, and the presence of high levels of reactive oxygen species (ROS) together with the reduction of apoptosis by antioxidants have suggested that ROS are relevant triggers of ICD (Cordeiro et al., 2019; Eshkar-Oren et al., 2015; Kaltcheva et al., 2016; Salas-Vidal et al., 1998; Schnabel et al., 2006). Macrophages have been detected in interdigital regions during ICD (Hopkinson-Woolley et al., 1994), however, it is not clear whether the recruitment of macrophages to the interdigital regions is a phenomenon concerted with the initiation of apoptosis or promoted by chemotactic signals secreted by apoptotic cells. In the present study we found that macrophages are positioned in the interdigital tissue before apoptosis initiation in most mesenchymal cells and produce ROS in association with phagocytosis.

## RESULTS

### Macrophages reside in the interdigits before abundant apoptosis initiation in this region

Apoptosis in the interdigits of developing mouse limbs correlates with the presence of professional macrophages (F4/80 ^+^ cells) (Hopkinson-Woolley et al., 1994; Wood et al., 2000). We dissected mouse embryo limbs before (S7 and S8) and during (S9) the onset of ICD. Although at S7 F4/80 ^+^ and TUNEL^+^ apoptotic cells were detected throughout the limb, they did not appear to be associated with ICD. Unexpectedly, we observed F4/80 ^+^ cells in the interdigits when very few TUNEL^+^ apoptotic bodies were detected in this region (S8; Fig. 1A); nearly all macrophages allocate in interdigits during this stage, since similar number of them were detected at the S9 and S9+ stages. Interestingly, however, macrophages at S8 stage showed a smaller size than those detected at later stages when apoptosis becomes abundant (Fig. 1B), suggesting that engulfment had not initiated. In the mouse, ICD initiates from distal and proximal interdigital areas, from which extends to the whole interdigit (Hernández-Martínez et al., 2009; Salas-Vidal et al., 2001). However, F4/80 ^+^ cells at S8 were already found populating the whole putative interdigital region (Fig. 1A; see also Fig. S1A), in contrast with the progressive pattern of ICD. Thus, it is apparent that macrophages prematurely define the interdigital area where ICD is going to occur.

**Figure 1.**
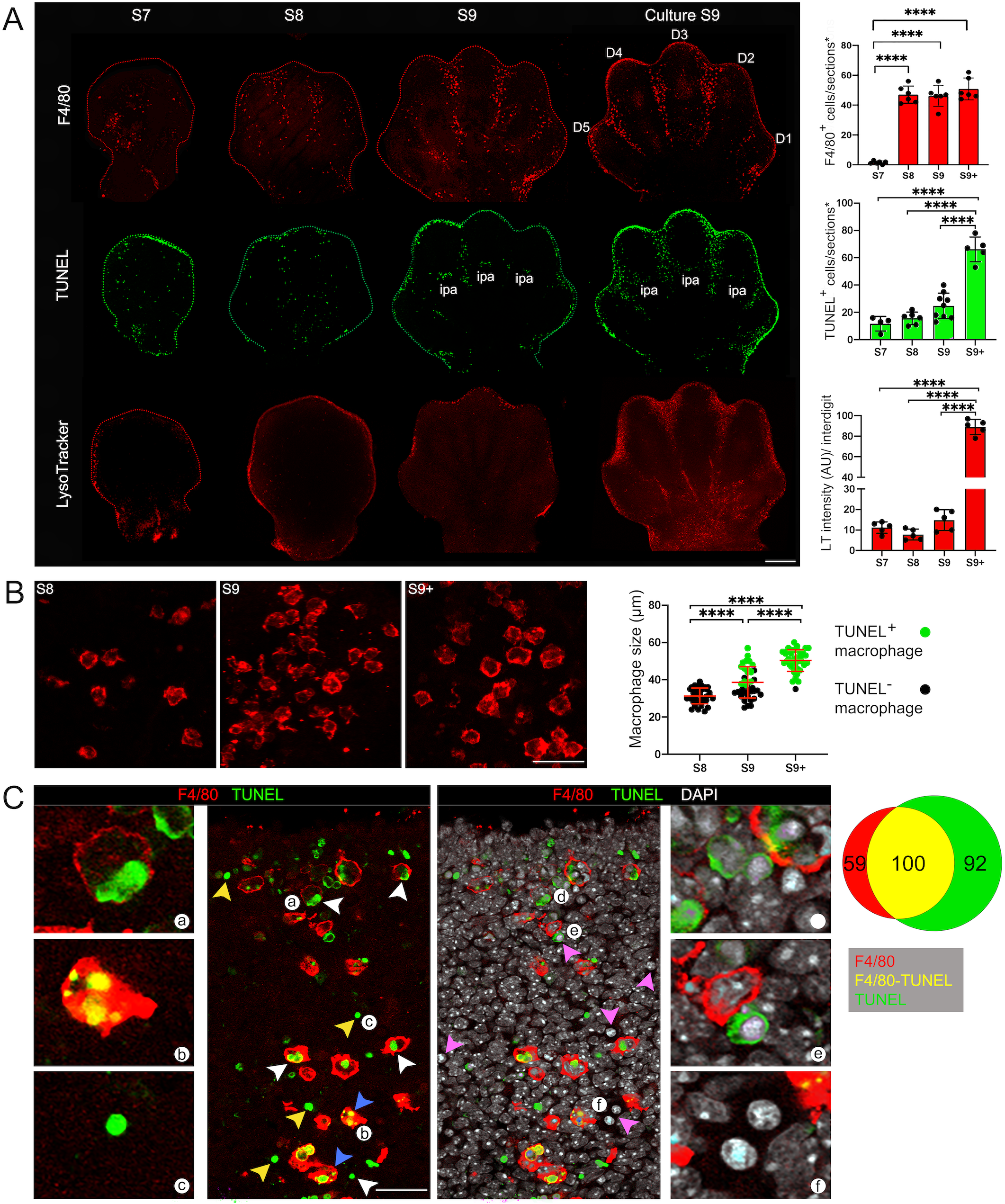
Macrophage distribution and phagocytic activity before and during ICD. (A) Macrophages (F4/80), apoptosis (TUNEL) and lysosomal activity (LysoTracker) in limbs at different developmental stages (S7-S9) and in cultured S9 limbs (scale bar, 500 µm). F4/80 and TUNEL were determined in sections containing most of the interdigital region, whereas LysoTracker staining was done on whole mount limbs (see details in Materials and Methods); images shown are the stack of 10-12 optical sections. Quantification of each marker is shown in graphs; for LysoTracker, the fluorescence intensity was quantified in the stack of optical sections (10-12) of each central interdigit of individual limbs (n=3), and the numbers of TUNEL^+^ and F4/80 ^+^ cells in graphs correspond to the positive cells found within 3 sections of each central interdigit of individual limbs (n=3) displaying the complete interdigital area (sections*; see details in Materials and Methods). D1-D5, digit identity; ipa, interphalangeal articulation. (B) Macrophage appearance at different developmental stages (S8-S9+; scale bar, 100 µm). Estimation of macrophage size is shown in graph (n=4). (C) Macrophages (F4/80), apoptotic cells (TUNEL) and condensed chromatin (DAPI) in S9+ limbs (scale bar, 100 µm). White arrowheads, engulfed apoptotic cells; blue arrowheads, engulfed apoptotic cells with a fragmented nuclei; yellow arrowheads, non-engulfed apoptotic cells. Lowercase letters (a-f) indicate cells shown in high magnifications panels. The proportion of cells with each marker is shown in the Venn diagram (3-4 intedigits analyzed).

As previously reported (Hopkinson-Woolley et al., 1994), macrophages residing in the interdigits showed limited migration towards apoptotic cells. Accordingly, F4/80 ^+^ cells were not detected in the developing interphalangeal articulations (ipa), where apoptosis was observed and are relatively close to macrophages in interdigits (Fig. 1A; see also Fig. 3C). In addition, about 47 ± 5% (n=4) of all apoptotic bodies with condensed chromatin (pink arrowheads) or positive for TUNEL (yellow arrowheads) were not associated with macrophages at the initiation of ICD (S9; Fig. 1C). Consistent with a poor attraction of macrophages by apoptotic cells, pharmacological caspase inhibition prevented apoptosis but did not affect macrophage presence in interdigits and, as expected, all macrophages in this condition were of small size (Fig. 2). In conclusion, the recruitment of macrophages to interdigital regions is not linked to apoptosis initiation and, rather, these two processes coincide during ICD.

**Figure 2.**
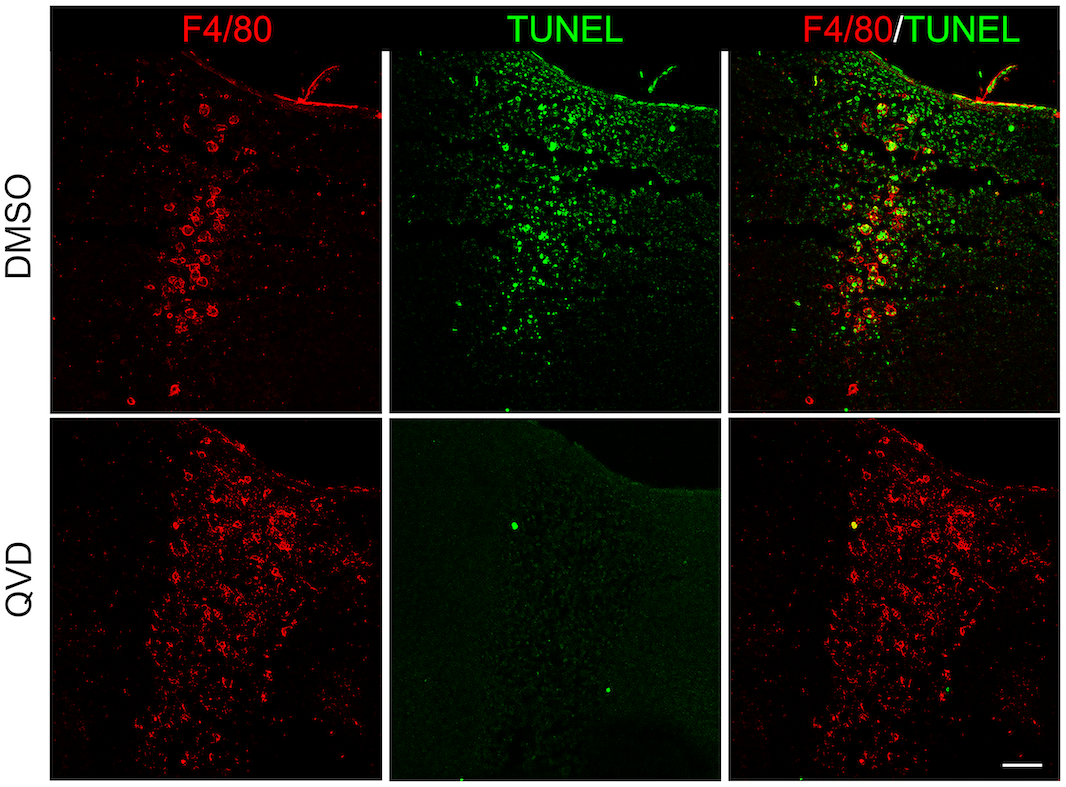
Macrophages in interdigital regions after apoptosis inhibition. S9 limbs were cultured for 8 h in the absence (DMSO) or presence of the pan-caspase inhibitor QVD. After culture, F4/80 (macrophages) and TUNEL (apoptotic cells) were determined in sections of treated limbs. Images of representative samples are shown (scale bar, 100 µm).

**Figure 3.**
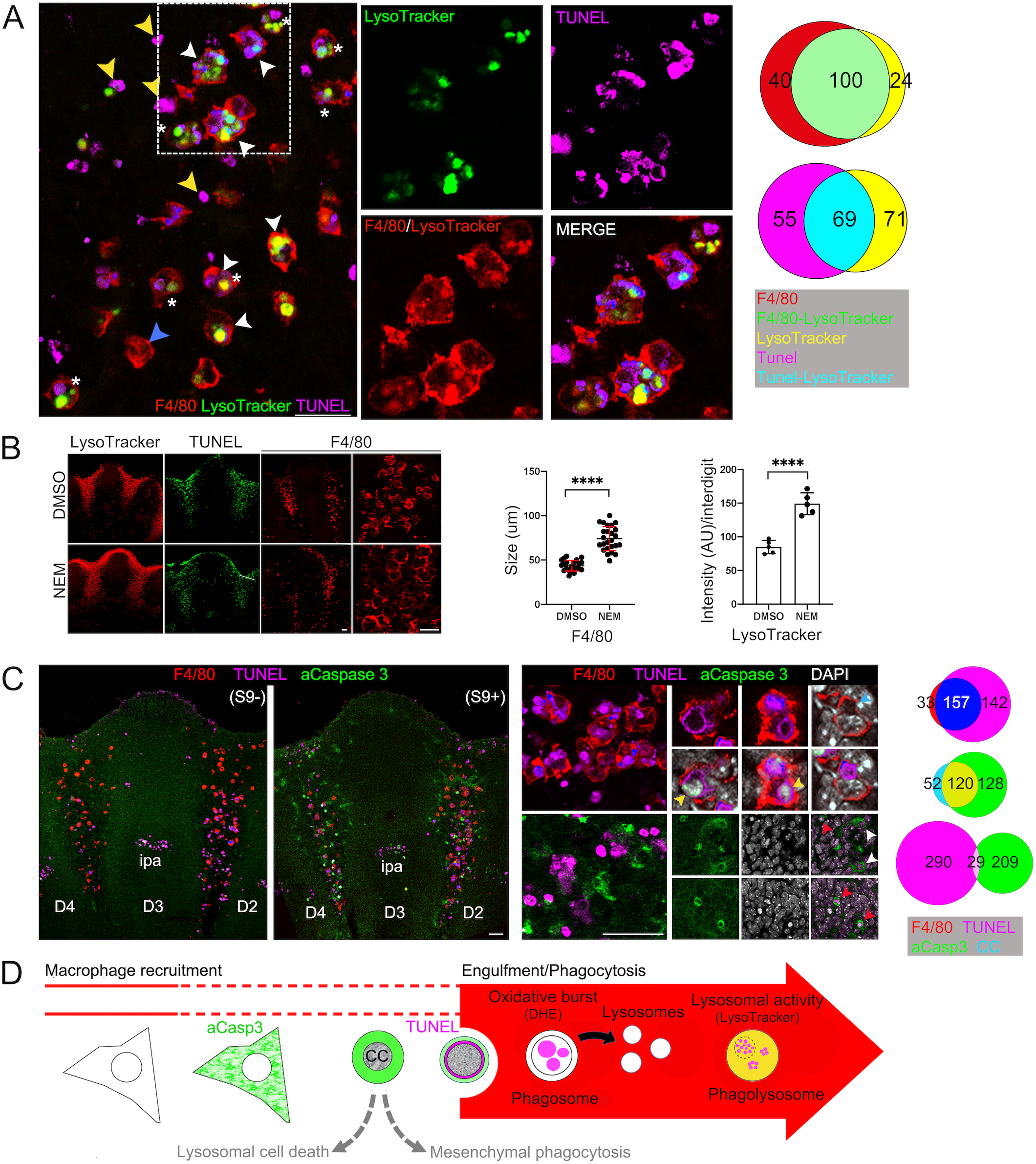
Engulfment and phagocytosis at the initiation of interdigital apoptosis. (A) Active lysosomes (LysoTracker) in interdigital macrophages (F4/80) during interdigital apoptosis (TUNEL) in limbs at S9. The association between the three markers analyzed is shown in the Venn diagram. White arrowheads, phagocytizing macrophages; blue arrowheads, non-phagocytizing macrophages; yellow arrowheads, non-macrophage associated apoptotic cells; asterisks, macrophages with phagosomes and phagolysosomes. (B) Macrophages (F4/80), apoptosis (TUNEL) and lysosomal activity (LysoTracker) in S9 limbs after induction of phagocytosis by NEM. Estimation of phagocytosis by macrophage size and by lysosomal activity (LysoTracker staining) is shown in graphs (n=3). (C) F4/80, TUNEL, aCasp3 and condensed chromatin in limbs before (S9) and after (S9+) distal LysoTracker detection. Magnifications of combinations of markers are shown in the right panels. Yellow arrowheads, nucleus with condensed chromatin surrounded by TUNEL signal; blue arrowheads, nucleus with non-condensed chromatin covered by TUNEL signal; red arrowheads, nucleus with condensed chromatin surrounded by aCasp3 signal; white arrowheads, nucleus with non-condensed chromatin surrounded by aCasp3 signal. Relevant associations between two markers are shown in Venn diagrams. (D) Schematic model of apoptosis and phagocytosis during ICD. Scale bars, 100 µm.

### High lysosomal activity correlates with phagocytosis but not with apoptosis during ICD

Although macrophages were found allocated before abundant ICD, phagocytosis activation must initiate shortly after apoptotic cells emerge. Phagocytosis starts when macrophages engulf apoptotic cells resulting in phagosomes that complete the internalization of apoptotic bodies. Phagosomes fuse with lysosomes and generate the phagolysosomes, characterized by their low intravesicular pH and in which a variety of hydrolytic enzymes “cleared” the engulfed content (Fairn and Grinstein, 2012; Stuart and Ezekowitz, 2005; Vernon and Tang, 2013). LysoTracker staining detects phagocyte degrading activity during ICD (Hernández-Martínez and Covarrubias, 2011). Accordingly, substantial LysoTracker staining correlated with the stages (S9 and S9+) at which a significant amount of apoptotic cells start to emerge and a high proportion of macrophages were of large size (47 ± 3 and 98 ± 6%, respectively; Fig. 1A,B). As expected, most of the LysoTracker ^+^ cells were also F4/80 ^+^ cells (80 ± 5%; Fig. 3A, white arrowheads; see also supplementary Fig. S1B) but, in agreement with the uncoupling of macrophage recruitment from apoptosis initiation, many F4/80 ^+^ cells located in interdigits were LysoTracker ^−^ and of small size (Fig. 3A, blue arrowheads; see also supplementary Fig. S1A). Interestingly, single F4/80 ^+^ cells were found with two types of large vesicles: those containing TUNEL ^+^ apoptotic bodies (putative phagosomes) and LysoTracker ^+^ vacuoles (putative phagolysosomes) (Fig. 3A, asterisks); this observation suggests that a single macrophage can asynchronously engulf several apoptotic cells. In addition, there were also LysoTracker ^+^/ F4/80 ^−^ cells, some found in the distal interdigital area at the initiation of ICD (S8, supplementary Fig. S1A) and others emerging throughout the degeneration process (S9+, supplementary Fig. S1B, green arrowheads). Therefore, although autophagic/lysosomal cell death or mesenchymal phagocytosis mechanisms might activate during ICD (LysoTracker ^+^/ F4/80 ^−^ cells), most lysosomal activity observed in interdigital regions is related to macrophage-mediated phagocytosis. Supporting this conclusion, treatment of limbs once apoptosis had initiated (i.e., S9) with NEM, a nonspecific compound that promotes PS exposure on the cell surface by irreversibly inhibiting the PS-flippase (Kuypers et al., 1996; Shiratsuchi et al., 1998), did not change the distribution of F4/80 ^+^ cells but, as expected, did increase engulfment (macrophage of larger size) and lysosomal activity (LysoTracker staining), exclusively at the interdigital regions (Fig. 3B).

It is apparent that macrophages initiate engulfment as apoptotic cells emerge in S9 limbs. To identify when apoptotic cells gain sufficient “eat-me” signals to be engulfed, apoptosis and engulfment were characterized in more detail in limbs before and after distal LysoTracker staining (S9 and S9+, respectively; see Material and Methods), representing the period at which ICD is initiating. In limbs before distal LysoTracker staining (Fig. 3C, S9; see Material and Methods), most TUNEL^+^ cells were restricted to proximal regions of interdigits, and after distal LysoTracker staining (Fig. 3C, S9+), TUNEL^+^ cells were also detected in distal regions (Fig. 3C, low magnification images). Interestingly, after detection of distal LysoTracker staining, cells with active caspase 3 (aCasp3 ^+^ cells) were distributed within the same areas as TUNEL^+^ cells but both makers were not coincident, indicating that they detect a different stage of the same apoptotic process (Fig. 3C, S9+). Notably, the distribution pattern of F4/80 ^+^ cells did not correlate with that of the TUNEL marker, particularly evident before phagocytosis (Fig. 3C, S9), and very few F4/80 ^+^/aCasp3 ^+^ cells were found (supplementary Fig. S2A). Once phagocytosis had initiated (Fig. 3C, S9+), nearly all macrophages (i.e., F4/80 ^+^/LysoTracker ^+^ cells) were positive for TUNEL and some had fragmented nuclei (Fig. 3A,C). Generally, perinuclear TUNEL staining was observed surrounding condensed chromatin (Fig. 3C, yellow arrowheads), and when the staining covered the complete nucleus, condensed chromatin was not evident (Fig. 3C, blue arrowheads); this pattern was observed within macrophages (Fig. 3C), but similarly occurred in apoptotic cells not associated with macrophages (supplementary Fig. S2B, yellow and blue arrowheads). Suggesting that activation of caspase 3 and chromatin condensation anticipates the DNA fragmentation detected by TUNEL along the apoptotic process, aCasp3 ^+^ cells had a rounded nucleus with non-condensed chromatin and some also a rounded cytoplasm with condensed chromatin, and most were negative for TUNEL (Fig. 3C, white and red arrowheads, respectively). Therefore, it is apparent that the great majority of cells dying in the interdigital region are executing a stereotyped apoptotic pathway, and many, but not all, are engulfed by macrophages and initiate phagocytosis at the DNA degradation phase (Fig. 3D).

### Inhibition of engulfment/phagocytosis does not prevent ICD

Pan-inhibitors of PI3K such as LY294002 can block phagocytosis by inhibiting PI3K class I and class III, enzymes involved in phagosome maturation (Bohdanowicz et al., 2010; Schlam et al., 2015; Thi and Reiner, 2012). Accordingly, the typical high lysosomal activity (LysoTracker ^+^) in interdigits of S9 limbs was abolished after 8 h of treatment with LY294002 in culture (Fig. 4A; see also (Hernández-Martínez et al., 2009); same result was obtained with another PI3K pan-inhibitor (PI-103; supplementary Fig. S3). However, treatment of S9 limbs with LY294002 also resulted in a significant reduction in F4/80 ^+^ cells and only few of them were found in interdigital regions of limbs after culture (Fig. 4A); most of these remaining macrophages had not initiated engulfment as determined by their size. This significant reduction in macrophages is likely due to the known role of PI3K class I signaling pathway in macrophage survival (Chang et al., 2009; Koh et al., 1998). Accordingly, inhibition of PI3K activity increased the number of TUNEL^+^ cells detected in interdigits (Fig. 4A), though added apoptotic cells could also derive from the expected lack of clearance of apoptotic bodies in macrophages.

**Figure 4.**
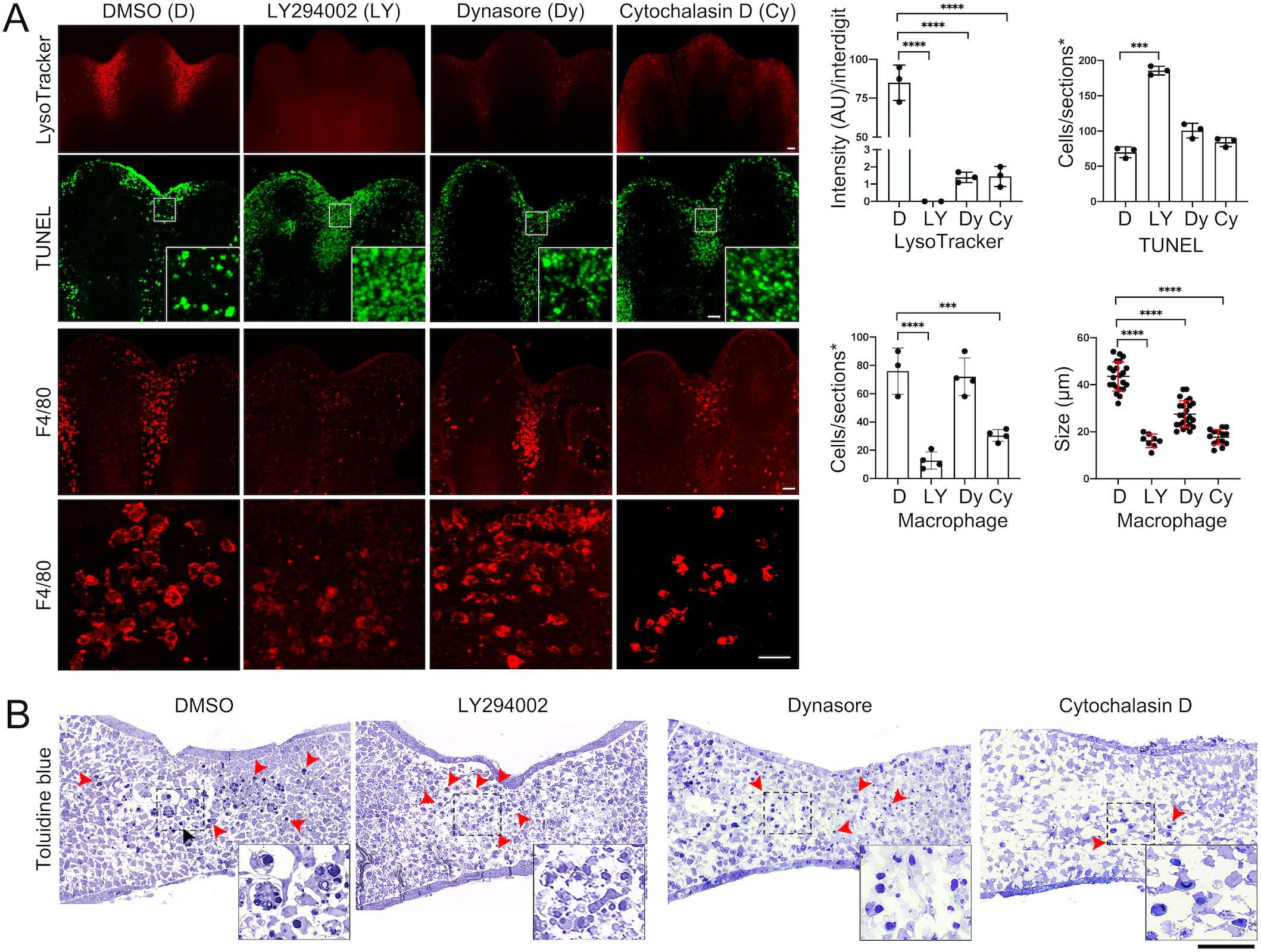
Phagocytosis and apoptosis after engulfment inhibition. Engulfment was inhibited by culturing S9 limbs for 8 h in the presence of LY294002, Dynasore and cytochalasin D. (A) Representative images and quantification of lysosomal activity (LysoTracker, whole mount determination), apoptosis (TUNEL), and of macrophage number and size (F4/80) in interdigits after limb culture are shown (n=3; scale bar, 100 µm). The numbers of TUNEL^+^ and F4/80 ^+^ cells in graphs correspond to the positive cells found within 3 sections of each central interdigit of individual limbs (n=3) displaying the complete interdigital area (sections*; see details in Materials and Methods). (B) Semithin cross sections stained with toluidine blue of limbs after culture with the reagents indicated (scale bar, 100 µm); red arrowheads point condensed nuclei. Insets are magnification of the interdigital areas shown. Insets in images (A, TUNEL, and B) show the distribution of apoptotic bodies in interdigits before (DMSO) and after engulfment inhibition (LY294002, Dynasore, Cytochalasin D).

Dynamin-2, a ubiquitously expressed dynamin family member that mediates the scission of endocytic vesicles from the plasma membrane, is required for engulfment by macrophages through a mechanism involving also PI3K class III (Gold et al., 1999; Kinchen et al., 2008). In the developing limb, abundant Dynamin-2 is found in interdigital engulfing macrophages (supplementary Fig. S4). Accordingly, the dynamin inhibitor Dynasore (Macia et al., 2006) inhibited engulfment and the subsequent lysosomal degradation and, as for the LY294002, did not prevent apoptosis in S9 limbs after 8 h of treatment in culture (i.e., reduced LysoTracker staining; Fig. 4A). In these cases, F4/80 ^+^ cells remained in the interdigital regions and the number of TUNEL ^+^ bodies were similar as control samples (Fig. 4A). The inhibition of engulfment was also noticeable by the smaller macrophage size in comparison with control limbs, though in the case of limbs treated with Dynasore, they were larger than those treated with LY294002 (Fig. 4A,B). Similar results were obtained with Cytochalasin D which, by inducing the depolymerization of actin cytoskeleton, can also block engulfment (Fig. 4A,B). Of note is that, when phagocytosis was inhibited, apoptotic bodies showed a dispersed pattern, in contrast with the regular patchy pattern formed by groups of engulfed apoptotic bodies within macrophages (Fig. 4A,B). Therefore, by blocking engulfment and/or phagosome maturation during ICD, detection of phagocytosis (e.g., LysoTracker staining) can be dissociated from the direct detection of apoptosis (e.g., TUNEL, aCasp3).

### **β**-galactosidase activity in interdigital regions detects active phagocytosis

β-galactosidase is a lysosomal enzyme, which high activity is associated with senescent cells (Kurz et al., 2000). β-galactosidase activity can easily be detected using X-gal or similar substrates (Lee et al., 2006). The optimal pH for β-galactosidase activity is pH4, but when the activity is very high, it can be detected at pH6, the condition for the so-called senescence-associated β-galactosidase (SA-β-galactosidase; (Krishna et al., 1999). β-galactosidase activity at pH6 has been detected in the interdigital regions of limbs and this evidence has been used to claim the presence of senescent cells during development (Lorda-Diez et al., 2015b; Muñoz-Espín et al., 2013; Storer et al., 2013). S9 and S10 mouse limbs were stained for β-galactosidase activity at pH4 and pH6 and, as expected, staining was observed in interdigital regions in both conditions but, with a stronger signal at pH4 than at pH6 (Fig. 5A). This β-galactosidase activity was reduced by the inhibition of engulfment and lysosomal degradation on cultured S9 mouse limbs using LY294002 or Dynasore (Fig. 5B). These data are in agreement with the high lysosomal activity in macrophages after engulfment and do not support the presence of senescent cells during interdigital tissue regression.

**Figure 5.**
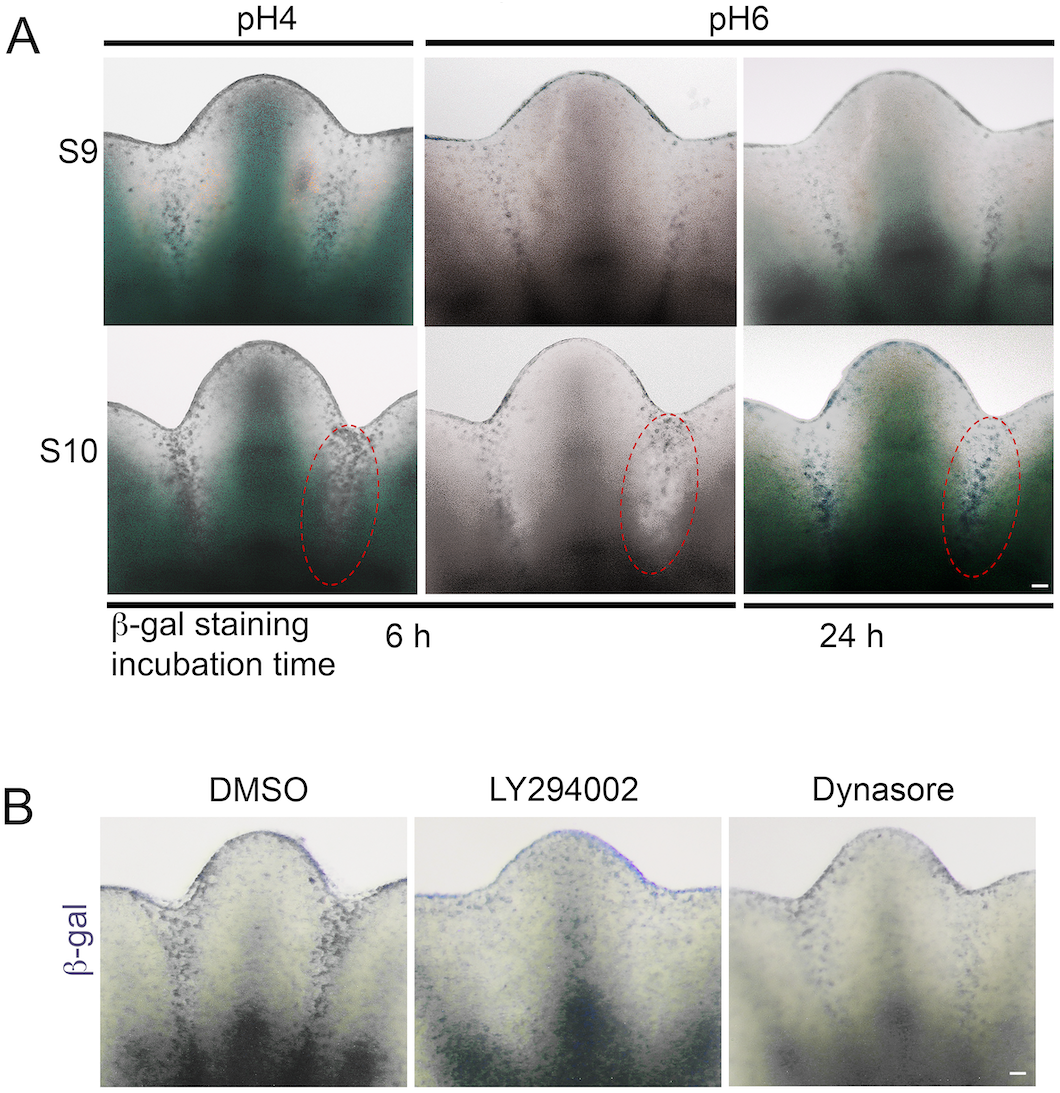
β-galactosidase activity in interdigits of developing limbs. (A) S9 and S10 limbs were stained for β-galactosidase activity at pH4 and pH6, and incubated for 6 or 24 hours. Note the weak signal at pH 6 that increased when incubation was extended for 24 h. (B) β-galactosidase activity after engulfment inhibition with LY294002 and Dynasore. Staining was performed after culture of S9 limbs with the reagents indicated for 8 h. Scale bars, 100 µm.

### Reactive oxygen species levels correlate with macrophage activity during ICD

Previously we reported that areas where abundant cell death is occurring in the developing mouse embryo correlate with regions showing high ROS levels (Salas-Vidal et al., 1998; Schnabel et al., 2006). In addition, these studies showed that antioxidants reduce apoptotic activity in interdigital regions (Salas-Vidal et al., 1998; Schnabel et al., 2006). However, although some cells with high ROS levels were identified as macrophages (Salas-Vidal et al., 1998), the source of ROS was not determined. ROS during ICD could be generated intrinsically from cells undergoing apoptosis or extrinsically from other cell types such as macrophages. To differentiate between these possibilities, we treated for 8 h S9 mouse limbs with either inhibitors of engulfment (LY294002 and Dynasore) or a generic antioxidant (TEMPOL) and determined ROS levels (Dihydroethidium) and lysosomal activity (LysoTracker). Inhibition of engulfment markedly decreased ROS levels (Fig. 6A, Dihydroethidium) and, interestingly, reducing ROS levels with TEMPOL did not affect lysosomal activity (Fig. 6A, LysoTracker). These results together suggest that ROS generation is a downstream event in phagosomes after engulfment and not coupled with the increase in lysosomal function (Fig. 3D). The NADPH oxidase inhibitor VAS2870 produced a similar effect as TEMPOL, indicating that macrophages generate ROS through the activation of NADPH oxidases (Fig. 6A; see Discussion). Macrophage number and engulfment was not altered by reducing ROS levels (TEMPOL, VAS2870) but, in agreement with previous observations (Salas-Vidal et al., 1998; Schnabel et al., 2006), fewer cells with fragmented DNA were detected (TUNEL^+^; Fig. 6B), suggesting that ROS contribute to DNA degradation in macrophages.

**Figure 6.**
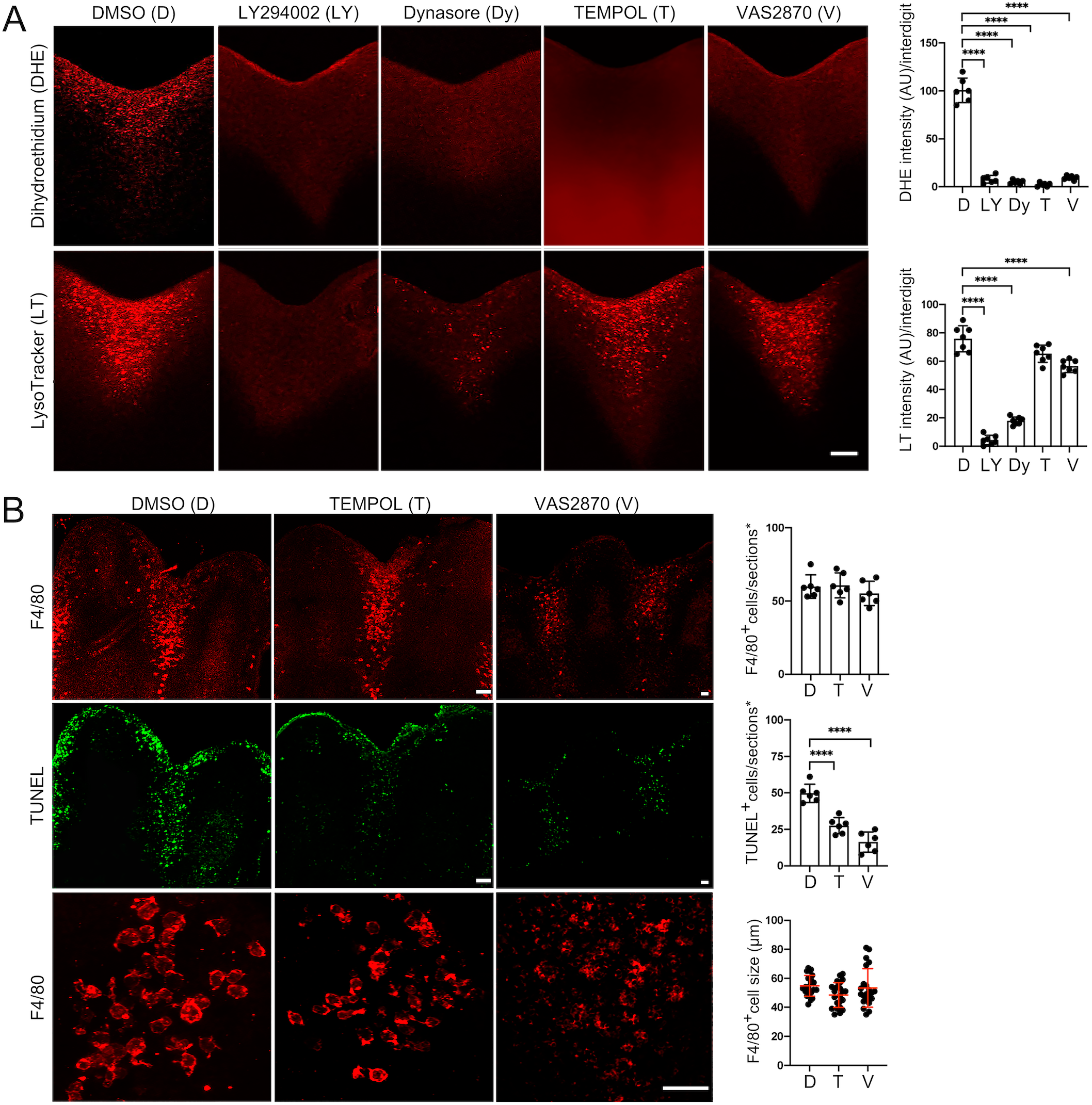
Phagocytosis and generation of ROS in association with ICD. (A) ROS levels (DHE) and lysosomal activity (LysoTracker) in S9 limbs treated (8 h in culture) with engulfment/phagocytosis inhibitors (LY294002, Dynasore) and antioxidants (TEMPOL, VAS2870). Representative images and quantification of ROS levels (DHE) and lysosomal activity (LysoTracker) are shown; DHE and LysoTracker levels were determined from the fluorescence intensity quantified in the stack of optical sections of each central interdigit of individual limbs (n=3). (B) Macrophages (F4/80 ^+^) and apoptotic cells (TUNEL^+^) after treatment with antioxidants for 8 h in culture. Representative images and quantification of macrophage number and size and of apoptotic cells are shown (n=3). The numbers of TUNEL^+^ and F4/80 ^+^ cells in graphs correspond to the positive cells found within 3 sections of each central interdigit of individual limbs (n=3) displaying the complete interdigital area (sections*; see details in Materials and Methods).

It has been proposed that blood vessels, which are abundant in the interdigits, regulate cell death by contributing to ROS generation (Eshkar-Oren et al., 2015). To test whether this mechanism involves macrophages, we determined their presence when blood vessel degeneration was induced using a specific Vascular Endothelial Growth factor Receptor inhibitor (VEGFR2i-III). As expected, adding VEGFR2i-III to the limb culture medium for 8 h induced endothelial cell death (supplementary Fig. S5) and caused a significant degeneration of the blood vessel network in the interdigital regions (Fig. 7A, CD31^+^). This condition did not alter the number of F4/80 ^+^ cells, and engulfment (Fig. 7A, F4/80 ^+^/TUNEL^+^) and lysosomal activity (Fig. 7B, LysoTracker) were not inhibited. Interestingly, however, the VEGFR2i-III inhibitor caused a marked reduction in ROS levels (Fig. 7B, Dihydroethidium). These data do not discard the contribution of blood flux to ROS generation as previously reported (Eshkar-Oren et al., 2015), but suggest that endothelial cells of blood vessels provide factors that possibly activate signaling pathways (e.g. those coupled to G protein receptors or mediated by MAP kinases) that are known to prime NADPH oxidases in immune cells (e.g., by phosphorylation of the p47^phox^ subunit; Nguyen et al., 2017) and, thus, determine ROS generation in the interdigital regions.

**Figure 7.**
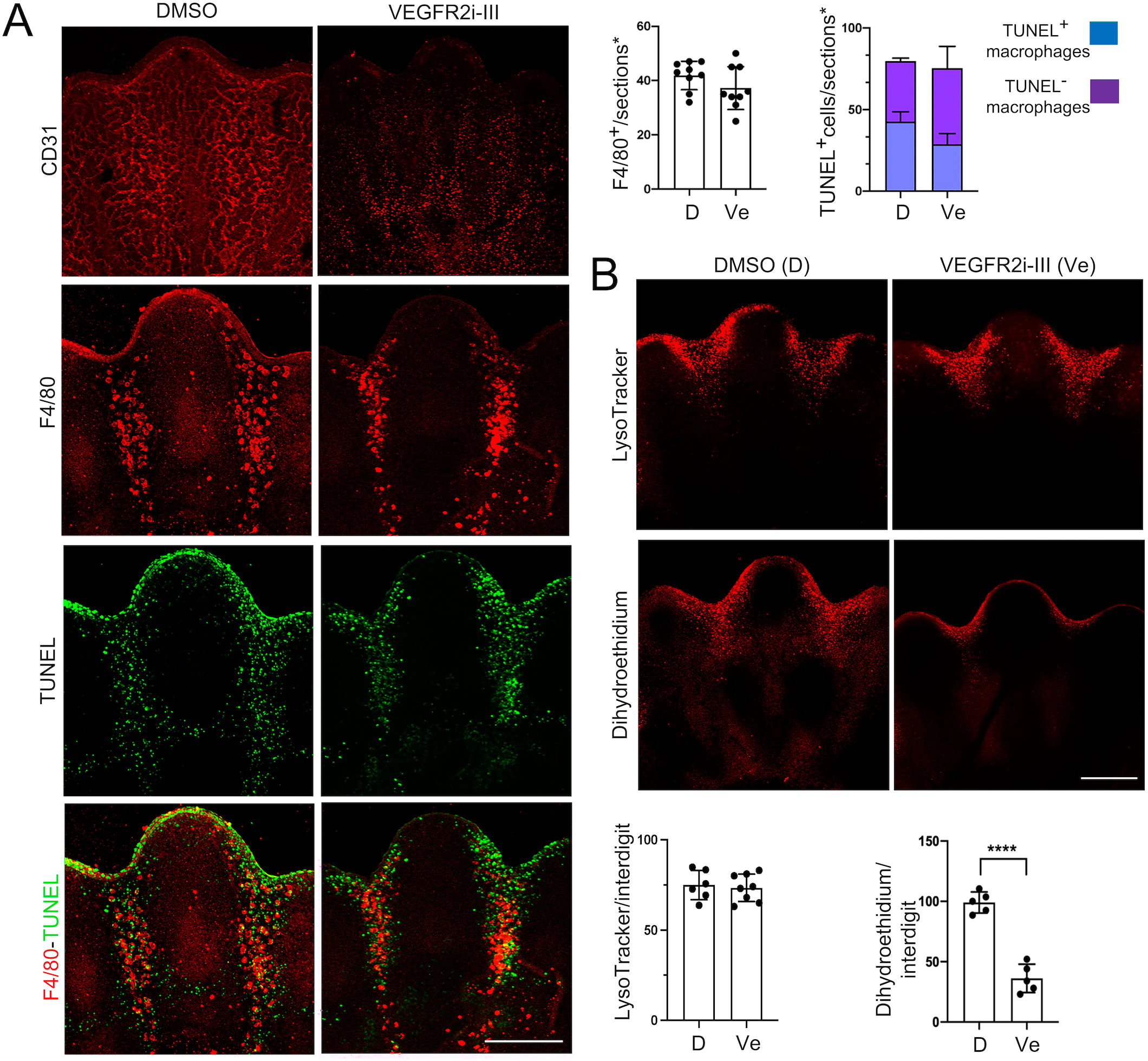
Effect of blood vessel integrity on macrophages, engulfment, lysosomal activity and ROS levels in the interdigital regions. Determinations were done in S9 limbs after 8 h treatment in culture with DMSO (vehicle) or with the VEGFR2 inhibitor VEGFR2i-III. (A) Representative images of central interdigits of limbs stained in whole mount for CD31 (blood vessels), and of limb sections stained for F4/80 (macrophages) or TUNEL (apoptotic cells) are shown (scale bars, 100 µm). The numbers of TUNEL^+^ and F4/80 ^+^ cells in graphs correspond to the positive cells found within 3 sections of each central interdigit of individual limbs (n=3) displaying the complete interdigital area (sections*; see details in Materials and Methods). (B) Representative images of central interdigits of limbs stained in whole mount with LysoTracker (lysosomal activity) or with dehydroethidium (DHE; ROS levels) are shown (scale bars, 100 µm). Graphs show the relative DHE and LysoTracker levels in each central interdigit determined from the fluorescence intensity in the stack of optical sections of individual limbs (n=3).

## DISCUSSION

Macrophages in the mouse embryo come from two different lineages: one that originates in the yolk-sac and the descendants of hematopoietic stem cells (HSCs), first located in the aorta-gonad-mesonephros and the fetal liver before reaching their final destination in the bone marrow (Perdiguero and Geissmann, 2016). Since most macrophages from the yolk sac distribute throughout the mouse embryo up to E11.5 dpc (named resident macrophages; Stremmel et al., 2018), it is likely that the significant number of macrophages in interdigits originate from circulating HSC-derived monocytes. Under this view, macrophages should reach the interdigital regions through the angiogenic signals that promote blood vessel invasion (Eshkar-Oren et al., 2015) and the still unknown signals that allow the extravasation into the interdigital tissue. Here we show that macrophages allocate in limb as digits develop but independent of the onset of ICD. Our observations contrast with the classical model in which it is expected that macrophages reach apoptotic regions following the “find-me”/“eat-me” signals produced by apoptotic cells (Lemke, 2019; Medina and Ravichandran, 2016).

### Recruitment of macrophages to interdigital regions is not linked to apoptosis

Macrophages have been found in developing embryos in association with the occurrence of apoptosis (Henson and Hume, 2006; Wood and Martin, 2017). In particular, macrophages are abundant in the interdigital regions of mouse developing limbs where a significant number of apoptotic cells are found (Hopkinson-Woolley et al., 1994). The apparent chronological coincidence of apoptosis emergence and macrophage allocation in different embryo regions is in accord with the recruitment of macrophages by apoptotic cells. However, as shown here, macrophages distribute in interdigital limb regions before significant apoptosis occurs and, in addition, macrophage motility seems to be restricted as macrophage distribution was not coincident with the progressive pattern of apoptosis emergence. This latter observation is in line with a previous report showing limited macrophage migration at early but not at late developmental limb stages (Hopkinson-Woolley et al., 1994).

The early allocation of macrophages to interdigital regions opens the possibility that they are involved in inducing apoptosis, as shown for endothelial cells in the rodent developing eye (Lobov et al., 2005) and for the natural death of Purkinje and motor neurons (Marı n-Teva et al., 2004; Sedel et al., 2004). However, it has been shown that apoptotic cells remain in interdigits of mice lacking PU.1, a master transcription factor required for the development of macrophages in almost all tissues, including those present in developing limbs (Wood et al., 2000). Notably, because the typical engulfment by macrophages is not occurring in this condition, apoptotic cells/bodies, instead of showing a patchy pattern, are dispersed within the interdigital region. We observed this same pattern when macrophages were almost completely depleted by the treatment of limbs with PI3K inhibitors or engulfment reduced by the dynamin inhibitor. Therefore, interdigital macrophages do not seem to contribute to the initiation of apoptosis but could speed the degradation of apoptotic cells (see below).

### Interdigital phagocytosis by macrophages is linked to high lysosomal activity

Several dyes used to detect apoptosis such as nile blue, neutral red, acridine orange (far-red emission) and LysoTracker are lysosomotropic dyes that are retained in acidic compartments (Hernández-Martínez et al., 2014). Thus, cells undergoing processes characterized by high lysosomal activity such as phagocytosis and autophagy can be detected with these dyes. Here we show that in the interdigital regions of developing mouse limbs, and likely in other regions with abundant cell death, these dyes specifically detect macrophage-mediated phagocytosis. Thus, LysoTracker staining increased when phagocytosis was promoted (e.g., in the presence of NEM) and decreased or not detected when phagocytosis was reduced (e.g., in the presence of PI3K inhibitors and Dynasore) or in the absence of macrophages (e.g., interphalangeal regions). Notably, no apoptotic cell before engulfment (i.e., aCasp3 ^+^/TUNEL^−^, aCasp3^+^/TUNEL^+^) showed high lysosomal activity, and only few cells with high lysosomal activity were detected not associated with macrophages. Almost all TUNEL^+^ macrophages contained also LysoTracker ^+^ vacuoles. As expected if phagocytosis occurs soon after apoptosis initiation, degradation of apoptotic bodies, as determined by lysosomal activity, follows a similar pattern as apoptosis (this study and Hernández-Martínez et al., 2009; Salas-Vidal et al., 2001), which contrast with macrophage allocation. Our observations are consistent with apoptosis being the major form of PCD in the limb interdigital region, and only few cells might be carrying out an autophagic/lysosomal type of cell death.

Cellular senescence is also associated with an increase in lysosomal activity (Kurz et al., 2000). This property has allowed the detection of senescent cells with the lysosomal activity of β-galactosidase. However, as shown here, in the interdigital regions β-galactosidase activity is almost completely associated with phagocytosis. Therefore, the claim for senescent cells in mouse embryos requires the detection of additional markers. However to discriminate cellular senescence from phagocytosis it is important to consider that p21 and p16, proteins commonly associated with cellular senescence, also have key regulatory roles in macrophages (Hall et al., 2017; Rackov et al., 2016). Similarly, at least some components of the senescence-associated secretory phenotype (SASP) are pro-inflammatory cytokines secreted by macrophages (Behmoaras and Gil, 2020). Therefore, caution should be taken to not misidentify senescent cells by the high β-galactosidase activity and other common markers of these cells, as they are also characteristics of phagocytizing macrophages.

### ROS contribute to phagocytosis

The significant reduction in ROS levels when phagocytosis was inhibited is a strong evidence that the increase in ROS in the interdigits, likely due to the activity of NADPH oxidases, is part of the phagocytosis process. ROS are known factors needed by macrophages to eliminate ingested microorganisms through the so-called oxidative/respiratory burst mediated by NADPH oxidases (Moghadam et al., 2021). Therefore, we propose that an oxidative-burst-like phenomenon contributes to the elimination of apoptotic cells during development. To our knowledge, this is the first evidence of the occurrence of an oxidative burst in macrophages during development. In accordance with this conclusion, the fact that antioxidants reduce the number of cells positive for the TUNEL in situ-DNA fragmentation assay (Cordeiro et al., 2019; Salas-Vidal et al., 1998; Schnabel et al., 2006), without altering engulfment, should be interpreted as the contribution of ROS to DNA degradation within macrophages rather than to the initiation of apoptosis.

High lysosomal activity and ROS levels might be properties of macrophages that allow them rapid degradation of engulfed apoptotic cells. As shown here, both properties were not observed in interphalangeal areas which naturally lack macrophages. Furthermore, the delayed regression of the interdigital tissue of limbs of *PU.1* null embryos lacking macrophages could result, not only from the inefficient engulfment by mesenchymal cells, but as shown by Wood et al., (2000), also from delayed degradation of apoptotic cells, as compared with that carried out by macrophages. Therefore macrophages could speed interdigital tissue regression by promoting rapid engulfment and degradation of apoptotic cells.

### Concluding remarks

Many markers used to detect PCD during development are shared by other cellular process such as phagocytosis, autophagy and cellular senescence. Specifically, here we show that in the interdigital regions of developing mouse limbs, and likely in other regions with abundant cell death, markers of lysosomal activity and ROS detect events of phagocytosis. Therefore, studies focused on the regulation of cell death should use additional markers of apoptosis in order to distinguish the molecules involved in the regulation of cell death itself from those regulating macrophage functions. In particular, a recent study reporting the role of environmental oxygen in the evolution of ICD (Cordeiro et al., 2019) might be uncovering the role of macrophage/phagocytosis in the evolution of limb morphogenesis, as ICD was only evaluated by LysoTracker staining.

Non-canonical functions of macrophages during development have recently been revealed (Theret et al., 2019; Wood and Martin, 2017). Macrophages have been implicated in morphogenesis, remodeling extracellular matrix and as a source of trophic factors. The early allocation of macrophages in interdigital regions before apoptosis initiation suggest other functions for macrophages than clearance of apoptotic bodies. It is tempting to speculate that macrophages and apoptosis evolved as independent processes that converge in developing regions with high tissue remodeling activity.

## MATERIALS AND METHODS

### Animals

Pregnant mice of the CD-1 strain were sacrificed by cervical dislocation from E11.5 up to E13.5 (0.5 days post-coitum was the day a vaginal plug was found; E0.5). Forelimbs or hindlimbs were indistinctly dissected and classified according with the limb bud development staging system of Wanek et al., (1989). Generally, S7 and S8 limbs were collected at early and late E12.5, respectively, whereas S9 and S10 at early E13.5. S9 limbs were also classified by those staining (S9+) or not staining the distal interdigital area with LysoTracker (i.e., before and after initiation of interdigital phagocytosis, respectively). In this case, whole limbs were stained with LysoTracker Red (L7528, Invitrogen), which can be detected before and after fixation and slicing (see below). All procedures were approved by the Bioethics Committee of IBT, UNAM and are in compliance with international guidelines.

### Mouse limb culture

The limb culture protocol used was as described by Salas-Vidal et al. (Salas-Vidal et al., 1998). Briefly, up to three S9-S9+ limbs were cultured over one 0.45 µm pore Durapore membrane filter (SLGVR33RS, Millipore) floating on DMEM medium (12100046, Gibco) for 8 hours at 37°C, and 5% CO_2_ within a humidified chamber. Dimethyl sulfoxide (DMSO; D2650 Sigma-Aldrich,), 30µM QVD (ab029, Kamiya Biomedical Company), 50 µM LY294002 (L9908, Sigma-Aldrich), 10 nM PI-103 (528100, Calbiochem), 60 µM Dynasore (324410, Sigma-Aldrich), 10 µM N-ethylemaleimide (NEM; E3876, Sigma-Aldrich), 20 µM VEGFR2i-III inhibitor (676487, Calbiochem), 0.3 µM Cytochalasin D (C8273, Sigma-Aldrich), 15 µM VAS2870 (SML0273, Sigma-Aldrich), and 10 µg/ml TEMPOL (581500, Sigma-Aldrich) were added to the DMEM medium just before culture. Except for Cytochalasin D, the integrity of the limb remained after the 8 h treatment with all the inhibitors added to the culture medium. In agreement with a minimal toxicity for limb cells by the inhibitors used (except cytochalasin), a significant change in the very low number of apoptotic cells in digits was not observed, and, except for cytochalasin D, semithin sections stained with toluidine blue did not reveal major abnormalities out of the interdigital zone.

### Dye stainings and histochemistry

Staining with LysoTracker (Green DND-26, L7526, or Red DND-99, L7528, Invitrogen), was performed on whole fresh or cultured limbs by incubating them in 1 µM solution for 15 min and then rinsed twice in PBS for 5 min, both at 37 °C. Same procedure was used for staining with 1 µM Dihydroethidium (DHE; D11347, Invitrogen). Whole limbs were stained for β-galactosidase activity at pH4 and pH6 by incubating them, for 6 or 24 h at 37°C, in the β-galactosidase tissue stain base solution (BG-8-C, Merck-Millipore) with either X-gal (B4252, Sigma-Aldrich) or bluo-gal (15519-028, Invitrogen) substrates. Stained samples were photographed in the MZ6 LEICA stereoscope using an Olympus camera (C-5050 ZOOM). Toluidine blue staining was performed on 800 nm semithin limb cross sections covering the most distal interdigital region. Samples were visualized by confocal microscopy (FV1000, Olympus) or by light microscopy (BX51, Olympus).

### Immunofluorescence and TUNEL assays

Determinations of TUNEL and markers detected by immunofluorescence were done in limb longitudinal cryo-sections that included the middle dorso-ventral zone of central interdigits (D2-D3 and D3-D4). D2-D3 and D3-D4 interdigits were selected for quantifications because: (i) they have very similar size and form thus, the number of apoptotic cells found are comparable; (ii) 3-D reconstructions of the limb show that central interdigits can be found within the same 2-D plane, consequently, the whole corresponding interdigital areas can be found within the same limb sections. Mouse limbs, fresh or cultured, were fixed in 4% paraformaldehyde (PFA) for 2h at room temperature, cryoprotected in 30% sucrose, embedded O.C.T. compound for 5h and frozen in the cryostat chamber at -25°C ensuring the right limb position for slicing; when required, samples were stored at -70°C. Longitudinal sectioning was done parallel to digits D2 and D4. Usually 6/6 µm sections, which cover the whole interdigital volume, were collected from each limb. Among these sections at least 3 showed a thick apical epithelium, full length D2-D4 digits and complete interdigital area (D2-D3 and D3-D4). Limbs stained with LysoTracker Red at the onset of ICD allowed to confirm that sections were around the middle dorso-ventral interdigital zone. Only sections comprising these characteristics were selected for determinations of markers and quantifications (see below). Following these criteria, it was evident to note that D2-D3 interdigit was always at a slightly more advanced stage than D3-D4 interdigit (previously described as a posterior-anterior gradient of ICD; Salas-Vidal et al., 2001). Immunofluorescence determinations were done using a standard procedure. Dilutions of antibodies were: 1:100, rat anti-mouse F4/80 (MCA497, BIO-RAD); 1:50, rabbit anti-active caspase 3 (aCasp3; 9664S, Cell Signaling); 1:500, Alexa Fluor^TM^ 546 goat anti-rat IgG (A11081, Invitrogen); 1:1000, Alexa Fluor^TM^ 594 goat anti-rabbit IgG (A11012, Molecular Probes). Terminal deoxynucleotidyl transferase-mediated dUTP nick end labeling (TUNEL) assay was performed using the In Situ Cell Death Detection Kit (11684795910, Roche), commonly, after the immunofluorescence procedure. Samples were stained with DAPI (D1306, Invitrogen) by incubating them in a 0.5 µg/ml solution in PBS for 10 min, followed by two rinses with PBS. When LysoTracker stained limbs were used, confocal images were taken after slicing and DAPI staining. Then, F4/80 immunofluorescence and TUNEL were performed on same samples and confocal images taken again; both images were merged using DAPI-stained nuclei as reference. Similar procedure was used for the double detection of aCasp3 and F4/80. A whole-mount immunofluorescence procedure was used for the detection of CD31. In this case, limbs were fixed with 4% paraformaldehyde for 2 h, washed in PBS for 10 min, and then dehydrated by sequential incubations for 10 min in 25%, 50%, 75% and 100% methanol in PBS. Limbs were rehydrated by incubations for 10 min in PBS and then blocked for 2 h in the blocking solution (PBS containing 10% goat serum, 1% triton X-100). Finally, samples were incubated in a 1:25 dilution of anti-mouse CD31 antibody (550274, BD Pharmingen) in blocking solution overnight at 4°C, followed by an overnight incubation in a 1:100 dilution of CY3 conjugated anti-rat IgG antibody (612-104-120, Rockland). Samples were visualized by confocal microscopy (A1R+ Storm, Nikon, or FV1000, Olympus); all images were captured from the dorsal limb side.

### Quantification and statistics

Number of macrophages (F4/80 ^+^), apoptotic cells (TUNEL^+^ and aCasp3 ^+^) and phagocytic cells (LysoTracker ^+^) were counted within the D2-D3 and D3-D4 interdigits of 3 physical sections comprising nearly all interdigital region (at least 90%). Counting was done through the analysis of 10-12 serial confocal images of each of the 3 selected physical sections (indicated as sections* in figures); sections that did not display the complete interdigital area, commonly the most ventral and the most dorsal sections showing reduced number of apoptotic cells, were not used for quantifications. Counting was performed in at least 3 limbs (n), in which D2-D3 and D3-D4 interdigits were individually evaluated. For counting cells with two or more markers, colocalization was confirmed by analyzing z orthogonal views. The macrophage size was estimated by measuring the maximum length of individual F4/80 ^+^ cells in 2-D images, selected after examining all z-optical planes covering every macrophage counted. Interdigital lysosomal activity (LysoTracker staining) and ROS levels (DHE staining), in fresh or cultured whole limbs, were determined on z-stacks confocal images (10-12) by measuring the relative intensity of the interdigital region using the Image J software in accordance with Du et al. (Du et al., 2020). Determinations were done in at least 3 limbs (n) of each developmental stage or treatment in culture, and D2-D3 and D3-D4 interdigits were individually evaluated. When three or more groups were analyzed, one-way parametric ANOVA was applied followed by either, pairwise comparisons using the Tukeýs multiple comparison test (Fig. 1A,B), or comparisons against a control group using the Dunnett’s test (Fig. 3A,B; Fig. 4A,B). When comparing two groups, an unpaired t-test was performed (Fig. 2B). P < 0.0001 (****) and P <0.001 (***) were considered significant. Statistical analyses were done using the GraphPad Prism software version 8.0b (La Jolla, CA, US). Venn diagrams were made using the Venn tool of the PowerPoint software 2022.

## ACKNOWLEDGEMENTS

We thank Dr. Mariana Gutiérrez-Mariscal for her helpful support in data statistical analysis and preparation of graphs. We also thank our institute animal facility staff (MVZ Graciela Cabeza Pérez, Oswaldo López Gutiérrez, Sergio González Trujillo and MVZ Elizabeth Mata Moreno) for their technical assistance. Confocal images were taken at the Laboratorio Nacional de Microscopía Avanzada with Dr. Verónica Rojo’s assistance and in Microscopy Core Facility of the Instituto de Investigaciones Biomédicas, UNAM, with Dr. Miguel Tapia-Rodríguez’s assistance. This work was partially supported by grants from the Programa de Apoyo a Proyectos de Investigación e Innovación Tecnológica (PAPIIT) of the Dirección General de Asuntos del Personal Académico (DGAPA) of the Universidad Nacional Autónoma de México (IN214219).

## AUTHOR CONTRIBUTIONS

Conceptualization: L.C., R.H-M., D.H-G.; Methodology: R.H-M., D.H-G., C.G-M., O.C-N.; Formal analysis: L.C., R.H-M., D.H-G., C.G-M; Investigation: R.H-M., D.H-G., C.G-M., O.C-N.; Data curation: L.C., D.H-G., C.G-M.; Writing - original draft: L.C.; Writing - review & editing: L.C., R.H-M., D.H-G., C.G-M., O.C-N.; Visualization: D.H-G.; Supervision: L.C.; Project administration: L.C.; Funding acquisition: L.C..

## SUPPLEMENTARY INFORMATION

**Figure S1.** Macrophages and lysosomal activity in interdigital regions. At the initiation of ICD, cells positive for LysoTracker were mainly found in the distal area (green arrowheads) and not associated with macrophages (F4/80 ^+^ cells) (A), whereas at a more advanced stage (S9+), most cells stained with LysoTracker were macrophages, but still few LysoTracker ^+^ cells (green arrowheads) were found not associated with macrophages (B). Note that just before ICD onset, several LysoTracker ^+^ cells were located underneath the apical ectoderm (delimited by white dots), and macrophages did not show signs of active phagocytosis (i.e., Lysotracker ^−^, small size). Blue arrowheads, non-phagocytizing macrophages; white arrowheads, phagocytizing macrophages. Scale bars, 100 µm.

**Figure S2.** The association of active caspase 3, TUNEL and condensed chromatin with macrophages. (A) The great majority of cells positive for aCasp3 were not associated with macrophages. (B) In agreement with a major apoptotic pathway for cells dying in interdigits, the TUNEL signal in the nucleus of apoptotic cells not associated with macrophages evolved as the TUNEL signal in apoptotic cells within macrophages (Fig. 2C); that is, first the TUNEL signal surrounded the condensed chromatin (yellow arrowheads), and when the TUNEL signal covered the complete nucleus, condensed chromatin was not detected (blue arrowheads). Scale bar, 100 µm.

**Figure S3.** Effect of PI-103 on S9 limbs in culture. S9 limbs were cultured in the absence (DMSO) and presence of PI-103 (PI3K inhibitor) for 8 h. LysoTracker staining was done in whole mount limbs, whereas TUNEL was performed on limb slices. Observe that the effect of PI-103 on LysoTracker staining and TUNEL signal was the same as that observed for LY294002.

**Figure S4.** Detection of Dynamin-2 in interdigital macrophages. Colocalization of F4/80 and Dynamin-2 in macrophages of S9 limbs. Observe that Dynamin-2 (green) was detected in nearly all F4/80^+^ cells (red) showing large size (engulfing macrophages).

**Figure S5.** Effect of VEGFR2i-III on S9 limbs in culture. S9 limbs were cultured in the absence (DMSO) and presence of VEGFR2i-III (VEGFR2 inhibitor) for 8 h. CD31 immunodetection and TUNEL were performed on limb slices. Note the disaggregation of blood vessels and the emergence of endothelial apoptotic cells (CD31^+^/ TUNEL^+^; arrowheads) in the presence of VEGFR2i-III.

