## Supplementary data for "Macrophages allocate independent of apoptosis and produce reactive oxygen species during interdigital phagocytosis"

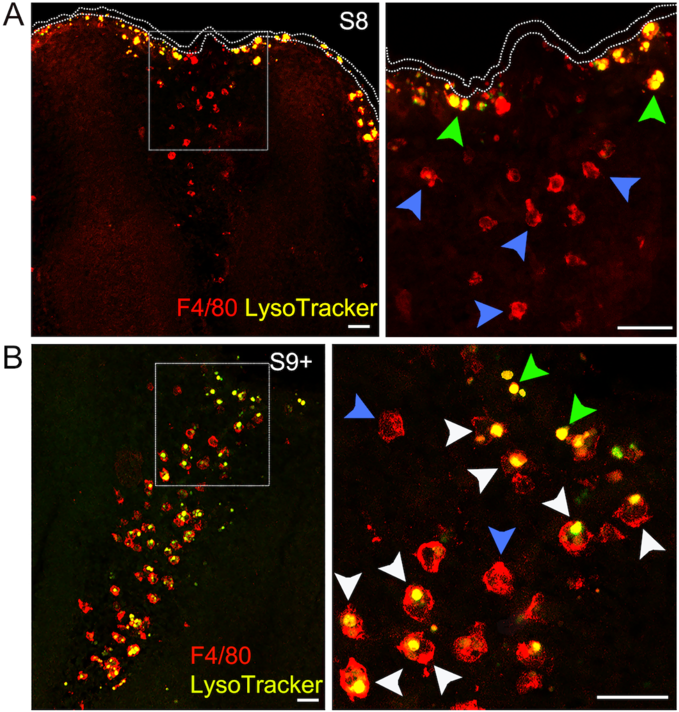


**Figure S1**

**Figure S1.** Macrophages and lysosomal activity in interdigital regions. At the initiation of ICD, cells positive for LysoTracker were mainly found in the distal area (green arrowheads) and not associated with macrophages (F4/80$^+$ cells) (A), whereas at a more advanced stage (S9+), most cells stained with LysoTracker were macrophages, but still few LysoTracker$^+$ cells (green arrowheads) were found not associated with macrophages (B). Note that just before ICD onset, several LysoTracker$^+$ cells were located underneath the apical ectoderm (delimited by white dots), and macrophages did not show signs of active phagocytosis (i.e., Lysotracker$^-$, small size). Blue arrowheads, non-phagocytizing macrophages; white arrowheads, phagocytizing macrophages. Scale bars, 100 µm.

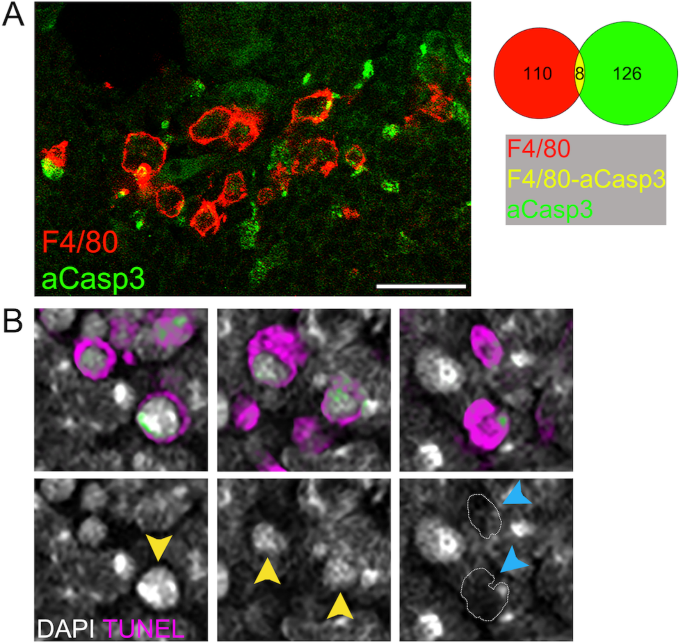


**Figure S2**

**Figure S2.** The association of active caspase 3, TUNEL and condensed chromatin with macrophages. (A) The great majority of cells positive for aCasp3 were not associated with macrophages. (B) In agreement with a major apoptotic pathway for cells dying in interdigits, the TUNEL signal in the nucleus of apoptotic cells not associated with macrophages evolved as the TUNEL signal in apoptotic cells within macrophages (Fig. 2C); that is, first the TUNEL signal surrounded the condensed chromatin (yellow arrowheads), and when the TUNEL signal covered the complete nucleus, condensed chromatin was not detected (blue arrowheads). Scale bar, 100 µm.

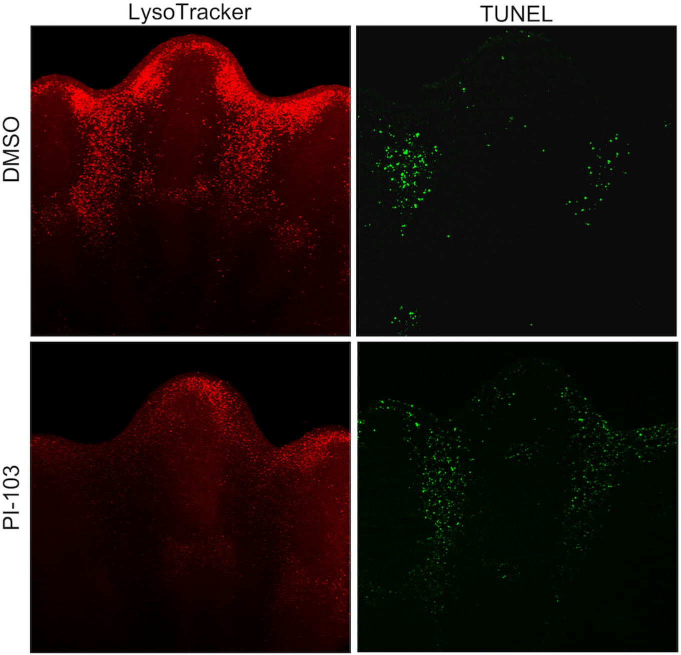


**Figure S3**

**Figure S3.** Effect of PI-103 on S9 limbs in culture. S9 limbs were cultured in the absence (DMSO) and presence of PI-103 (PI3K inhibitor) for 8 h. LysoTracker staining was done in whole mount limbs, whereas TUNEL was performed on limb slices. Observe that the effect of PI-103 on LysoTracker staining and TUNEL signal was the same as that observed for LY294002.

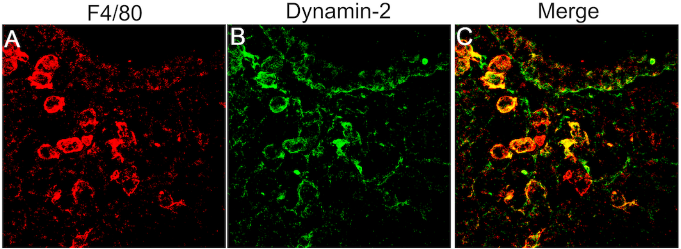


**Figure S4**

**Figure S4.** Detection of Dynamin-2 in interdigital macrophages. Colocalization of F4/80 and Dynamin-2 in macrophages of S9 limbs. Observe that Dynamin-2 (green) was detected in nearly all F4/80$^{+}$ cells (red) showing large size (engulfing macrophages).

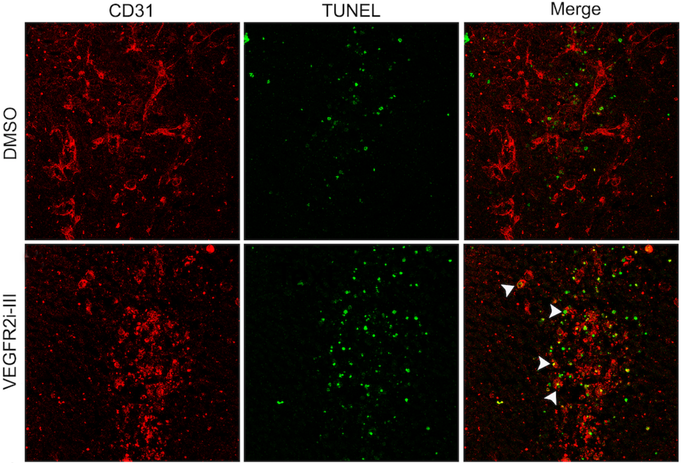


**Figure S5**

**Figure S5.** Effect of VEGFR2i-III on S9 limbs in culture. S9 limbs were cultured in the absence (DMSO) and presence of VEGFR2i-III (VEGFR2 inhibitor) for 8 h. CD31 immunodetection and TUNEL were performed on limb slices. Note the disaggregation of blood vessels and the emergence of endothelial apoptotic cells ($CD31^+$/ $TUNEL^+$; arrowheads) in the presence of VEGFR2i-III.
